# Energy and heterogeneity shape bird taxonomic and functional gamma-diversity patterns across landscapes in Finland

**DOI:** 10.64898/2026.04.13.717752

**Authors:** Jérémy Cours, Aleksi Lehikoinen, Daniel Burgas, Risto K. Heikkinen, Merja Elo, Martijn Versluijs, Rémi Duflot

## Abstract

**Aim:** Our aim was to study the effects of energy availability and landscape habitat heterogeneity on bird taxonomic and functional gamma-diversity and propose conservation guidelines based on the results.

**Location:** Southern and Central Finland

**Time Period:** 2009–2020

**Major Taxa Studied:** Birds

**Methods:** We derived biodiversity variables from bird monitoring line transects to assess the effects of latitude, longitude, and landscape composition, configuration, and heterogeneity at multiple spatial scales: 100, 500, 2,000, and 5,000 m. We tested the effects of these landscape metrics on the total community, bird ecological guilds (species richness and abundance), functional diversity, and overall species specialization index.

**Results:** We found clear evidence supporting a positive effect of energy (latitude and soil fertility) and habitat amount on bird abundances. Our results also revealed a northward increasing trend in functional diversity and species specialization. Habitat heterogeneity positively affected both bird abundance and species richness. Heterogeneity of land cover types was shown to promote abundances, while functional measure of landscape heterogeneity was positively connected to species richness. Land use with high anthropogenic activities, such as urban areas and cropland, negatively affected forest specialists and species sensitive to human activities.

**Main Conclusions:** Energy and habitat heterogeneity and amount are major mutually nonexclusive factors shaping bird communities in Finnish landscapes. Nonetheless, certain land use types favour some guilds while excluding others (for example, urbanized areas or cropland favouring open area species while excluding old-growth forest specialists), showing that biodiversity conservation is a matter of specialized landscapes. Furthermore, different measures of landscape heterogeneity demonstrated positive relationships with the studied bird guilds, highlighting the consistency of the species–heterogeneity relationship.

## 1. Introduction

As biodiversity confronts multiple pressures from anthropogenic activities, particularly land use change and intensification, with one million species currently threatened with extinction (IPBES, 2019), it is urgent to improve the understanding of the key ecological mechanisms involved in structuring species communities. At the landscape scale, three main mechanisms have been identified (Honkanen et al., 2010). First, the *species–energy relationship* describes how increasing solar energy with decreasing latitude (hereafter referred to as energy) is related to increasing species richness (Evans et al., 2005; Hawkins et al., 2003; Root, 1988; Wright, 1983). This could be due to several reasons. Higher energy availability could allow a higher number of individuals to co-exist in the regional species pool, indirectly increasing species richness (the *more-individual hypothesis*; Evans et al., 2005; Mönkkönen et al., 2006). Furthermore, higher energy can support larger populations, which may decrease species extinction rates and/or increase long-term speciation (Evans et al., 2005; Honkanen et al., 2010). Higher energy levels may also allow for the presence of higher trophic levels within community networks, resulting in higher species richness (Evans et al., 2005; Honkanen et al., 2010).

Second, habitat filtering has been suggested as another important factor that acts through habitat heterogeneity and habitat amount (Leibold et al., 2004). The *habitat heterogeneity hypothesis* states that habitat diversity in landscapes enhances gamma-diversity and primarily affects species richness through niche partitioning (Cours & Duflot, 2025; Duflot et al., 2022; Fahrig et al., 2011; Hurlbert, 2004; Tews et al., 2004). Landscape heterogeneity is composed of two dimensions: compositional (number of habitats) and configurational (spatial arrangement of habitats); both are supposed to interactively promote gamma-diversity (Duflot et al., 2022; Fahrig et al., 2011). Higher compositional heterogeneity translates into higher niche richness, while higher landscape configuration translates into higher habitat edge density. Such ecotones may be considered specific habitats with their own associated species that promote complementation processes (Dunning et al., 1992).

The *habitat amount hypothesis* (or *habitat area hypothesis*) states that an increase in habitat in a landscape leads to more species dependant on that habitat. This pattern can be attributed to the increase in the number of individuals with the increase in the habitat amount, facilitating higher species richness (the *more-individual hypothesis*; Fahrig, 2013; Honkanen et al., 2010; MacArthur & Wilson, 1967), and is related to the *species–energy* hypothesis, connecting the per-unit-area productivity with the total energy available (resources) to a particular group of species (Clarke & Gaston, 2006; Wright, 1983). In parallel, a larger habitat amount increases the colonization– extinction ratio, ultimately allowing for a higher species number to survive (Fahrig, 2013; Honkanen et al., 2010; MacArthur & Wilson, 1967).

Knowledge gaps persist regarding the hierarchy of these mechanisms, their additivity, and the direction of their effects. For instance, whether fragmentation (configurational heterogeneity) has positive or negative effects on biodiversity is still debatable (Valente et al., 2023). Furthermore, while the effects of certain factors (e.g. habitat heterogeneity) have long been identified (Duelli, 1997; Tews et al., 2004), the methodology for characterizing them needs to be improved. For example, landscape habitat heterogeneity has long been assessed through land cover typologies (e.g. diversity of land cover types), which are strongly anthropocentric and might not efficiently capture the habitat characteristics relevant to communities, leading to potentially spurious results (Fahrig et al., 2011; Valente et al., 2023). Additionally, land cover typologies do not provide information about the heterogeneity factors that drive species responses; for example, in forests, it is not clear whether variations in canopy openness or tree diameter matters for bird communities (Cours & Duflot, 2025). Thus, an improved understanding of these mechanisms is necessary to implement efficient measures for biodiversity conservation.

We investigate the additive and mutually nonexclusive effects of energy, landscape heterogeneity, and habitat amount on bird communities in Finnish forest landscapes. Four decades of monitoring have highlighted strong bird population decreases in Europe and North America, particularly among farmland species (Rigal et al., 2023; Rosenberg et al., 2019). A similar trend is noted in forest species, especially specialists associated with declining and changing habitats in the boreal zone (Betts et al., 2022; Cours et al., 2025; Virkkala, 2016). Birds are highly mobile organisms that may move across large distances. Therefore, they are sensitive to landscape habitat conditions and are relevant for examining landscape–biodiversity relationships (Cours & Duflot, 2025).

The additive and mutually nonexclusive effects of energy, landscape heterogeneity, especially when measured at fine resolution, and habitat amount on bird communities remain unclear. Bird community responses to landscape conditions have mostly been studied within protected areas (Brotons et al., 2003; Elo et al., 2012; Honkanen et al., 2010). It is important to extend bird community studies to all available landscapes in Finland and identify the main drivers of bird communities to better account for them in future conservation planning. Moreover, recent developments in species traits and specialization indices databases (Le Viol et al., 2012; Pearman et al., 2014; Tobias et al., 2022) provide opportunities for studying specific ecological guilds, community trait values, and functional diversity indices (Cours et al., 2025). For instance, Bae et al. (2018) found that taxonomic and functional indices inversely respond to energy availability.

In our study, we elucidate the relative importance of the three main mechanisms – energy, habitat heterogeneity, and habitat amount – driving species communities at the landscape scale. Specifically, we studied the gamma-diversity of bird communities across the southern and central Finnish forest production landscapes using the line transect data of the national bird breeding monitoring scheme from 2009 to 2020. We iteratively included a large array of metrics in generalized linear mixed models to test the effects and importance of each mechanism on the taxonomic diversity index (i.e. abundance and richness of all species and ecological guilds) and functional diversity index (i.e. functional diversity indices and community-weighted means). We hypothesized that (i) energy is the main positive factor for bird abundances (*more-individual hypothesis*), (ii) habitat heterogeneity is the main factor for bird species richness, and (iii) habitat amount is an important factor for the richness and abundance of bird species associated with that particular habitat (guilds).

## 2. Materials and methods

### 2.1. Bird gamma-diversity

We utilized observation data from the line transects of the Finnish monitoring of breeding birds (Virkkala & Lehikoinen, 2014). The line transect method is a one-visit census typically conducted from late May to late June, depending on geographical position. The transects are about 6 km in length and are systematically distributed within 25 × 25 km grids across Finland. We selected transects that were visited from 2009 to 2020 in the hemiboreal, southern, and central boreal zones in Finland (Fig. 1), matching the time slice with the availability of land cover data. We excluded the northern boreal region from our analyses because it has very different landscape settings (Fig. 1). Moreover, we removed transects overlapping national borders or the sea within a 10-km buffer (since we focused only on terrestrial landscape processes). Consequently, our analysis was conducted using a selection of 1,577 transect-year observations (337 transects unevenly visited every year from 2009 to 2020; average number of visits = 4.7; Fig. 1). Given the length of the line transects, we ensured that they provide a multihabitat, landscape-level measure of bird diversity (i.e. gamma-diversity). For each transect, we calculated the species richness and abundance (number of breeding pairs), accounting for differences in species detectability for the latter (Virkkala & Lehikoinen, 2014). Detailed methodology of the line transect surveys is provided in Supplementary Material S1.

**Fig. 1.**
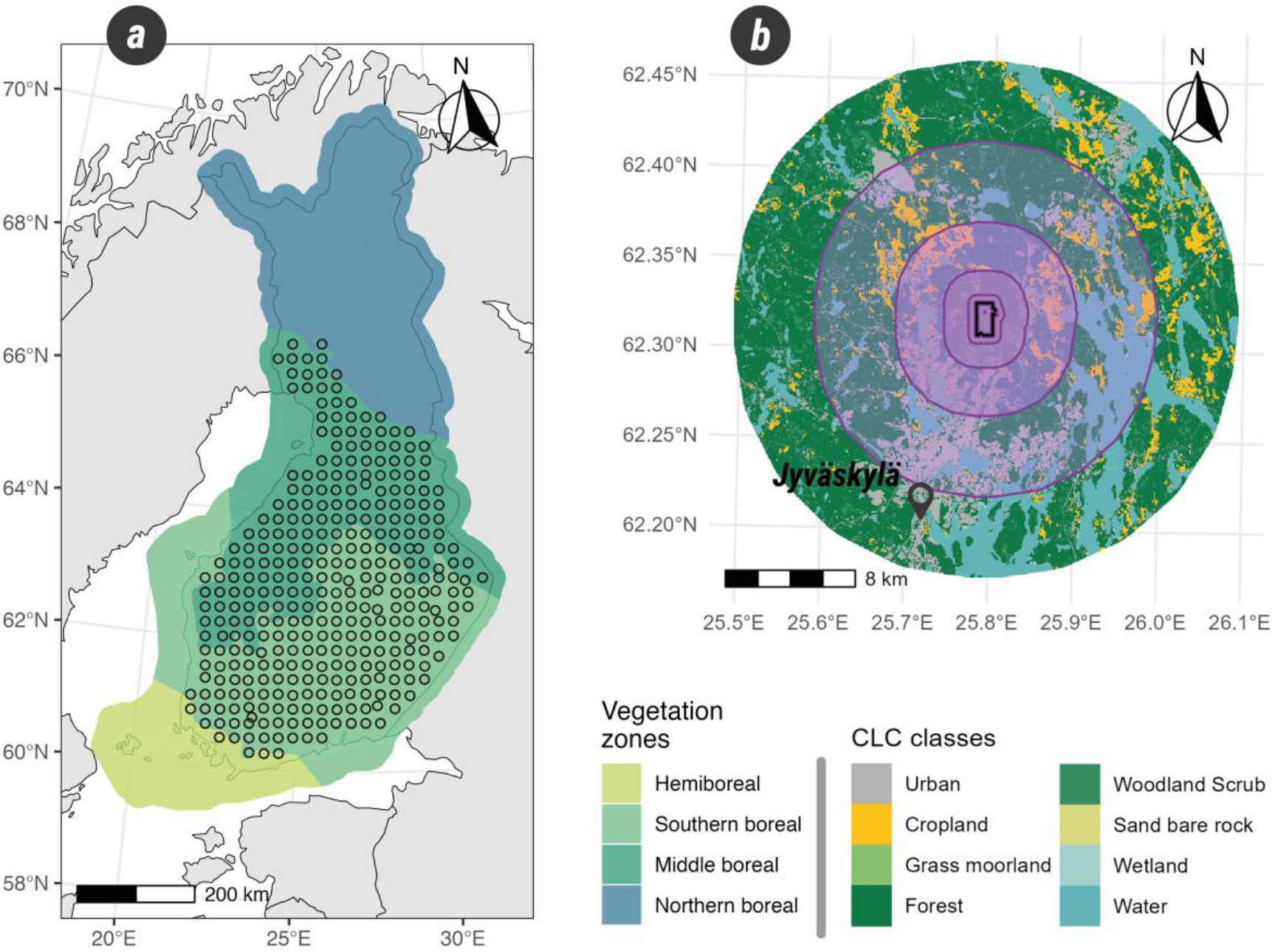
(a) Map of bird transect censuses selected for the landscape analysis (i.e. in the hemiboreal, southern, and middle boreal zones). (b) Example of a transect (black near-rectangular line) and the surrounding landscape next to the city of Jyväskylä, Finland. Transparent purple buffers define the landscape at different scales – 100, 500, 2,000, and 5,000 metres – around the transect.

As species characteristics often determine their response to environmental gradients, we opted for a functional diversity approach. We collected the conservation status from the Red List of Finnish Species (Lehikoinen et al., 2019) and extracted data on species, including average adult body mass and primary habitat and trophic niche information, from Tobias *et al*. (2022). In addition, we collated migration status (Howard et al., 2023), nest location (Pearman et al., 2014), old-growth forest specialization (Fraixedas et al., 2015; Mönkkönen et al., 2014), overall species specialization index (Le Viol et al., 2012), peak human tolerance index (Marjakangas et al., 2024), and species temperature indices (Lehikoinen et al., 2021).

We calculated the abundance and species richness of the overall communities (all species), as well as at guild-level — i.e. forest and old-growth forest specialists, and open-habitat species associated with grassland and shrubland. Within the forest species, we also calculated the abundance and richness of forest and resident and hole-nesting species. By selecting the species with the lowest human tolerance (lower quantile of observed species), we calculated the abundance of human-avoiding species. By using the red-list status of species as an ordinal gradient, along with their body mass, habitat preference, trophic niche, nest location, and migration strategy, we calculated the indices of the functional diversity of the overall communities (all species), functional richness divergence, and Rao’s Quadratic Entropy (RaoQ). We calculated the same functional indices for forest species using the same traits, replacing only the overall habitat preference with the specialization of old-growth forests. We noted the presence-absence of red-listed forest and red-listed open-habitat species. Finally, we calculated the community-weighted mean (CWM) of the specialization index for all species (Le Viol et al., 2012), which was estimated as the mean of the specialization value of each species and weighted by their abundance.

### 2.2. Landscape heterogeneity metrics

We utilized open-source data to characterize the landscapes around the bird line transects, matching bird observations to spatial data from the closest corresponding year and at multiple scales of 100 m, 500 m, 2 km, and 5 km. First, we harmonized the land cover classes of the Corine Land Cover (CLC) maps for 2006, 2012, and 2018 (see Table S1 for details on land cover class grouping for each year). Second, we obtained a more detailed description of forest structure and composition using the multi-source national forest inventory (MS-NFI), updated every two years from 2009 to 2019. The MS-NFI is an extrapolation of the field-based Finnish national forest inventory based on remote sensing data, mostly from Sentinel and Landsat (Mäkisara et al., 2022). The MS-NFI provides raster layers of mean canopy closure; tree diameter, height, and age; total tree volume; volume of main tree species (pine, spruce, birch, and other broadleaf); and site fertility categories. We resampled the CLC (resolution of 25 m in the 2006 map and 20 m in the maps for the following years) at the MS-NFI resolution (20 m in 2009 and 2011 and 16 m in the following years) and converted the continuous variables from the MS-NFI into discrete classes (see Table S2).

From these land cover maps, we derived regular landscape composition and configuration metrics and calculated the proportion of croplands, grass and moorland, urban areas, wetlands, and water bodies. The proportion of forests was evaluated by considering pixels classified as forest, woodland and scrubland, and wetland on the CLC map and those with a canopy closure > 20% to account for planted forests on peatlands and regenerating clear-cuts. Using the proportion of each of the CLC classes, we calculated the Shannon diversity index as an overall measure of landscape heterogeneity. In addition, we measured the proportion of mixed (broadleaf and coniferous trees < 80% of total tree volume), broadleaf (broadleaf trees > 80% of total tree volume), and old forests (age ≥ 100 years). Finally, we calculated several habitat configuration metrics: mean patch area, patch density, mean core area (with an edge width of one pixel), total core area, edge density, and mean perimeter–area ratio for the patches of the CLC categories and mixed, broadleaf, and old forests. To better represent the total variability of terrestrial seminatural habitats (excluding urban areas, croplands, and water bodies), we created vegetation types based on both CLC and MS-NFI maps, ranging from bare ground habitats to more detailed forest types (see Table S3). We then calculated the Shannon diversity index based on the proportion of these vegetation types, as well as the abovementioned configuration metrics.

We also opted for a functional heterogeneity approach to measure habitat variability based on structural variables from the MS-NFI. We allocated an ID to each pixel based on the unique combinations of CLC categories, ordinal classes from MS-NFI, and seminatural vegetation types (Fig. S1). These IDs were further used to represent habitat species, with the aggregated information corresponding to their traits. Subsequently, we treated each landscape buffer as a sampled area, in which each pixel represents an individual of each habitat species. We used the noncorrelated traits (*r* Pearson < 0.6) associated with the forest structure (canopy closure, tree diameter, age, total volume, and broadleaf volume; see Fig. S2) to calculate landscape functional richness (FRic), RaoQ, and divergence (FDiv; Fig. S1). We calculated them for the entire landscape (i.e. all habitats) and for the seminatural habitats only (as defined above). For forested habitats only, we calculated the CWM and functional dispersion (FDis) of canopy closure, tree diameter, age, total volume, broadleaf volume, and site fertility. We inversed the site fertility scale compared to the data provided so that a higher number corresponding to class fertility denotes greater fertility (1 = unproductive lands, 10 = herb-rich forests; Mäkisara et al., 2022).

For the analysis, we selected landscape metrics with correlation < 0.8, keeping only one of two or more correlated metrics (see Table S4 for the selected variables and their definitions; see Figs. S3–S10 for the correlation tables). We removed the proportion of grass and moorland since the values were relatively low (0%–9%) at the 500 m scale and did not vary much across landscapes.

### 2.3. Analysis

R software v.4.5.1 was used for our analysis (R Core Team, 2025). We found that landscape metrics were strongly correlated with latitude (e.g. thinner trees towards the north due to shorter growing season). Because we were interested in testing the effect of latitude as a proxy for available energy, in addition to specifically testing the effects of landscape heterogeneity and habitat amount independent from the latitude effect, we extracted the residuals from the loess relationships between each landscape metric and latitude and used them as predictors. We used generalized additive models (GAM; *mgcv* R-package) to evaluate the landscape effects on bird communities. Depending on the biodiversity variable, we used Poisson (for species richness), negative binomial (for abundances), Tweedie (if fitting better than the two first), or Gaussian (for functional indices and specialization indices) distribution, all including a logarithmic link function (see Fig. S11 for the mean and range values of biodiversity variables). We used binomial distribution (logit link function) for the presence-absence of red-listed species and added the logarithm of abundance as a fixed covariable in each richness model to estimate the effects of landscape metrics independently from the potential indirect effects of abundance. We added the logarithm of species richness in each functional richness model due to the large correlation between the two and the interaction term between latitude and longitude in all the models to control for spatial autocorrelation (see Figs. S12 and S13). We also added transect ID and sampling year as random factors and the log-transformed transect lengths as an offset to account for varying total transect lengths.

We structured our analysis in several stages. First, for each response variable, we selected the metrics best describing the proportion of primary land covers at the optimal buffer scale based on Akaike information criterion (AIC; i.e. proportions of forest, cropland, urban areas, wetland, and water bodies within a buffer of 100, 500, 2,000, or 5,000 metres). This set of variables remained fixed in subsequent stages. Next, we created simple models for each of the remaining landscape metrics, testing one metric and one buffer scale at a time, and ranked them based on AIC. In the third stage, for each biodiversity response variable, we performed a forward stepwise selection by successively adding the best landscape metrics based on simple models. We evaluated all metrics whose simple models had a negative ΔAIC compared to the model obtained in step 1, indicating an improved fit. When the inclusion of a metric variable did not improve the model fit (ΔAIC > −2), it was not retained in the analysis. We also tested the additive effects of the same metric at different buffer scales, provided that their variance inflation factors (VIF) were less than three (i.e. low collinearity across scales of the same landscape metric; Zuur et al., 2010). This resulted in a few cases in which the same metric at different scales was retained in the final model. This process resulted in the best multiscale model for each response variable.

## 3. Results

### 3.1. Effects of energy availability and broad biogeographic conditions on bird communities

Although we found an overall reduction in bird abundance and species richness northwards, except for abundances of human-avoiding species and birds of prey (positive) or the species richness of open-habitat species (no effects, Fig. 2), functional diversity, specialization, and the presence of red-listed forest species increased northwards (Figs. 2 & 3).

**Fig. 2.**
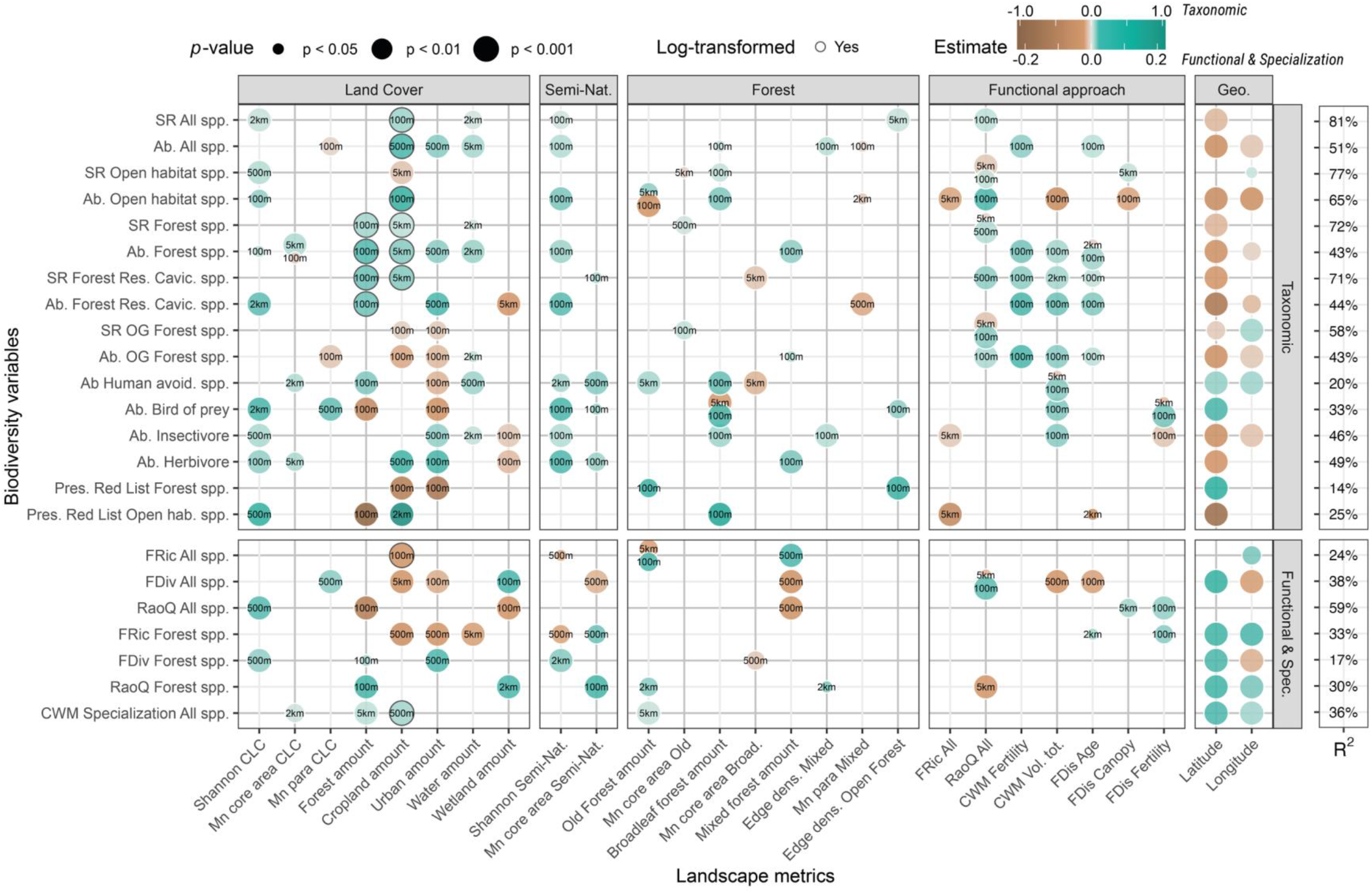
Graphical representation of the effects of significant landscape metrics on bird taxonomic diversity (top) and functional and specialization indices (bottom). Each row represents a model. The numbers in bubbles indicate the landscape scale at which the effects were detected. R-squared values denote the explained deviance in each model. To focus on the main relationships, we only display landscape metrics affecting more than two biodiversity variables (see Fig. S16 for full details).

**Fig. 3.**
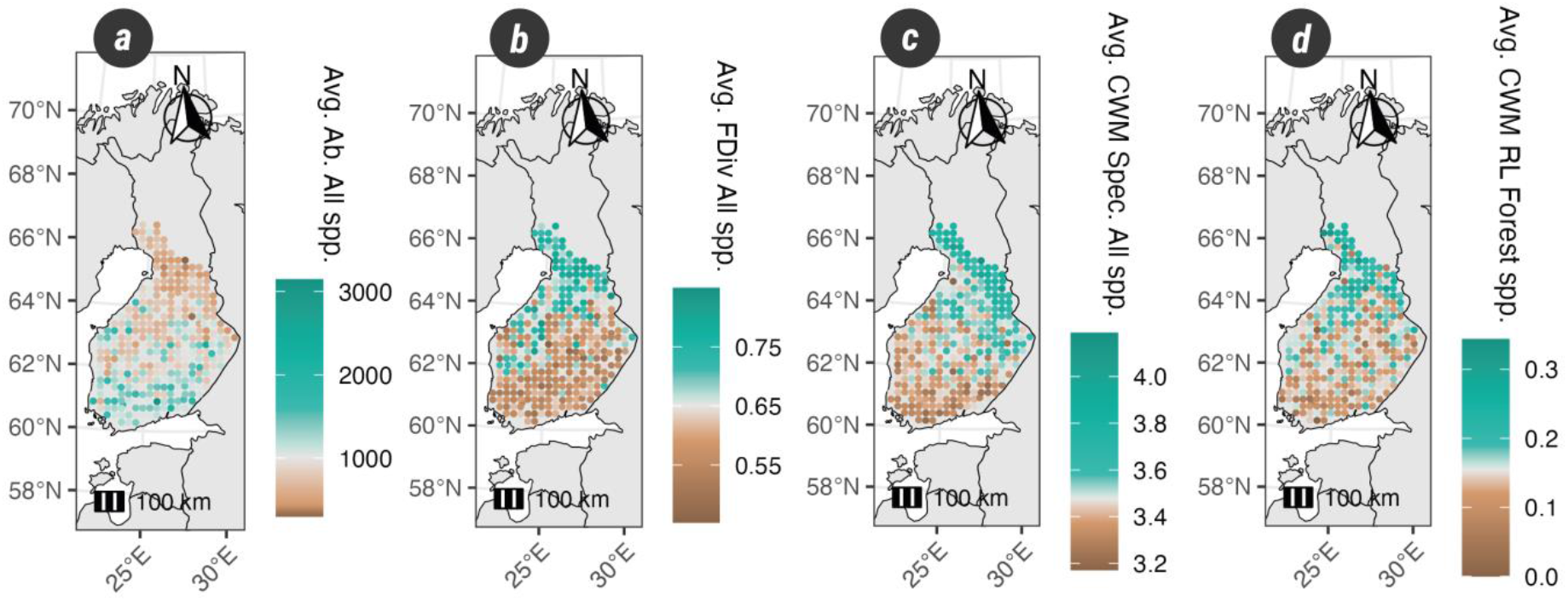
Values of biodiversity variables for each transect in Finnish hemiboreal, southern, and middle boreal zones (averaged across years): (a) overall abundance, (b) functional divergence, (c) CWM habitat specializations for all species, and (d) CWM Red List of forest species.

Soil fertility generally had positive effects on most of the taxonomic biodiversity variables at the 100 m scale. It increased the abundances of all species and of forest species, including the old-growth forest specialists and the abundance and species richness of forest resident hole-nesters (Fig. 2). Similarly, higher levels of total tree volume at the 100 m scale increased the abundances of the different forest guilds and the human-avoiding species, birds of prey, and insectivores (Fig. 2), but it had a negative effect on the functional divergence of all species.

### 3.2. Effects of landscape heterogeneity on bird communities

#### 3.2.1. Effects of compositional heterogeneity

Landscape heterogeneity displayed an overall positive effect on different biodiversity variables, but the scale of effect varied depending on the landscape metric and biodiversity variable (Fig. 2). We found that the Shannon diversity index of CLC was positively related to the species richness of all species at the 2 km scale and of open-habitat species at the 500 m scale (Fig. 2). Shannon diversity index of CLC promoted the abundances of forest resident hole-nesters (2 km scale), birds of prey (2 km scale), insectivores (500 m scale), and herbivores (100 m scale), the presence of red-listed open-habitat species (500 m scale), the functional diversity (richness [5 km scale] and RaoQ [500 m scale]) of all species, and the functional divergence of forest species (500 m scale; Fig. 2). The Shannon diversity index of seminatural vegetation types measured at 100 m had a positive effect on both the abundance and species richness of all species and the abundances of open-habitat species, forest species, including forest resident hole-nesters, and the three trophic guilds. It also increased the functional divergence of forest species (2 km scale) and the CWM of the specialization index (100 m scale), while it had a negative effect on the functional richness of forest species (500 m scale; Fig. 2).

Regarding the habitat functional approach, RaoQ of all habitats promoted most of the guild species richness, mostly at 100 and 500 m scales (Fig. 2), and the abundances of open-habitat species and old-growth forest specialists (100 m scale; Fig. 2). It also increased the functional divergence of all species (100 m scale), while it led to a decrease in the functional RaoQ of forest species at 5 km scale (Fig. 2). We also found many similar effects of the different functional dispersion indices within forested habitats. For example, tree age dispersion measured at the 100 m scale promoted the total abundance and the abundances of forest species, including the resident hole-nesters, and of old-growth forest specialists (Fig. 2). Canopy variations decreased the abundance of open-habitat species while positively affecting their species richness (Fig. 2). We noticed that variations in seminatural site fertility increased the functional RaoQ of all species and the functional richness of forest species (Fig. 2).

#### 3.2.2. Effects of configurational heterogeneity

We found that an increased mean core area of patches of different land covers or seminatural habitats benefitted several biodiversity variables (abundances of human-avoiding species [2 km scale], herbivores, and birds of prey; functional richness and RaoQ of forest species [500 and 100 m scales], and the CWM of the specialization index [2 km scale]; Fig. 2). Patch complexity, measured as the perimeter–area ratio (para), of land cover had negative effects on the abundance of all species, especially on old-growth forest specialists (100 m scale). In contrast, it benefitted the functional divergence of all species and the abundance of birds of prey at 500 m scale (Fig. 2).

Edge density between forest and open habitats promoted the species richness (5 km scale) of all species, abundance of birds of prey (100 m scale), and the presence of red-listed forest species (100 m scale; Fig. 2). Increasing the old forest core area benefitted the richness of forest and old-growth specialist species (at 500 and 100 m scales; Fig. 2). The mean core area of broadleaf forests negatively affected the abundance of human-avoiding species and the species richness of forest resident hole-nesters at the 5 km scale, as well as the functional divergence of forest species (500 m scale; Fig. 2). Mixed forest edge increased the abundances of all species and herbivorous species (100 m scale) and the functional RaoQ of forest species (Fig. 2). Mixed forest patch complexity negatively affected the abundance of all species (100 m scale), open-habitat species (2 km scale), and forest resident hole-nesters (500 m scale; Fig. 2).

### 3.3. Effects of habitat area on bird communities

Overall, forest proportion had positive effects on forest guilds at the 100 m scale, including their functional divergence and RaoQ (Fig. 2), but negative effects on the presence of red-listed open-habitat species (5 km scale) and birds of prey (100 m scale) guilds and on the RaoQ of all species (100 m scale; Figs. 2 and 4). Cropland and urban proportions had similar negative effects at the 100 m scale on old-growth forest specialists and on the presence of red-listed forest species, indicating less endangered species in areas with high proportions of these land covers (Fig. 2). Cropland and urban areas also significantly reduced the functional divergence of all species and the functional richness of forest species (Fig. 2).

**Fig. 4.**
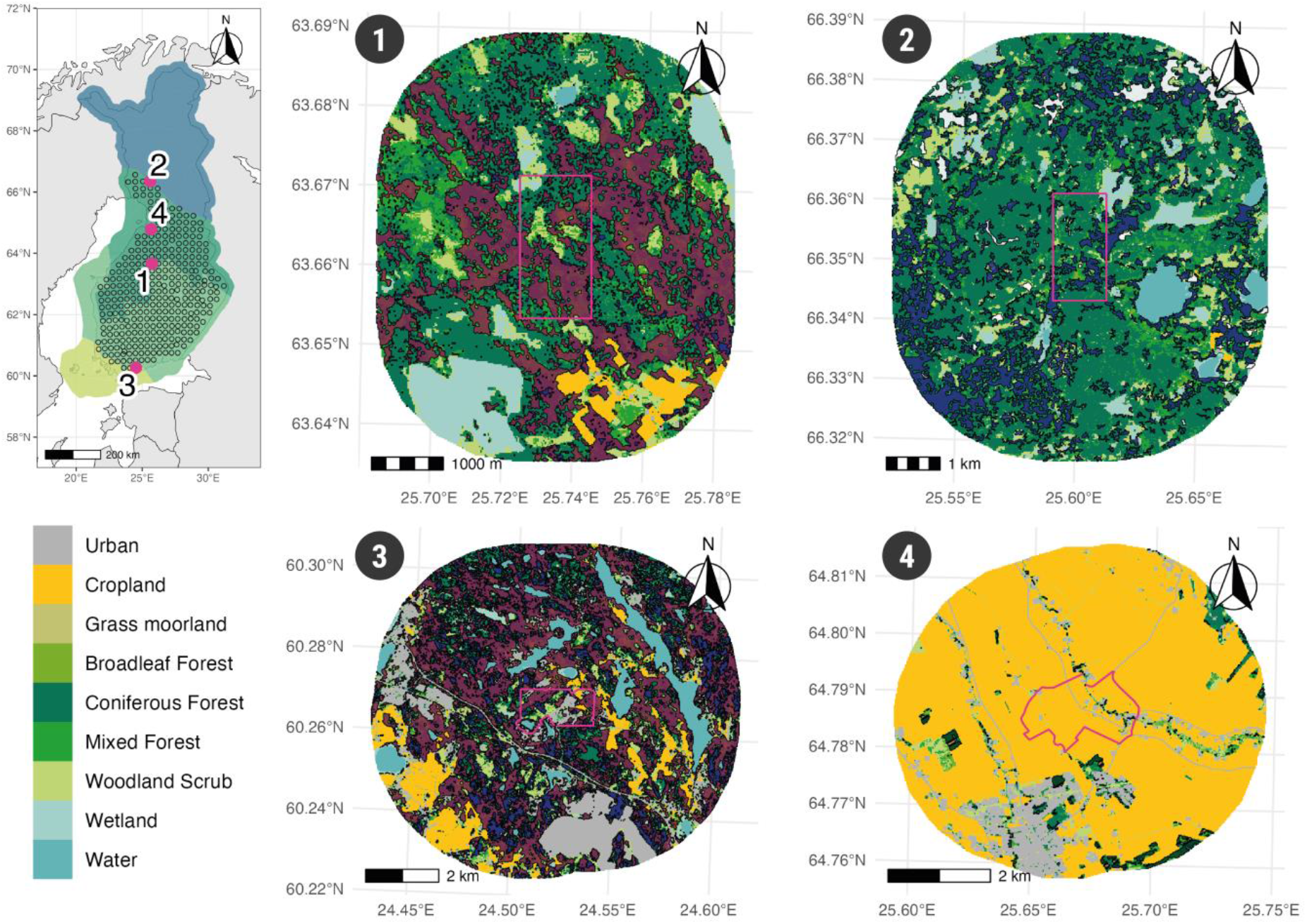
An example of four different landscapes suitable for four different bird guilds: (1) old-growth forest specialists, (2) red-listed forest species, (3) forest resident hole-nesters, and (4) open habitat bird species. Dark purple areas represent mature forests with a total wood volume > 200 m^3^.ha^-1^, mostly in (1) and (3). Dark blue areas represent old forests > 100 years, mostly in (2). White areas in (2) represent clear-cuts.

Cropland and urban areas promoted the abundance of all species (100 m and 2 km scales), forest species (5 km and 500 m scales), and herbivores (500 m and 100 m scales). Cropland proportion increased the richness of all species (100 m scale) and forest species (5 km scale), the presence of red-listed open-habitat species (2 km scale), and the CWM of specialization index (500 m scale). Croplands favoured the abundance of open-habitat species at 100 m scale but reduced their species richness at 5 km scale (Fig. 2). It also decreased the functional richness of all species. The proportion of urban areas at 100 m or 500 m scales led to a decrease in the functional divergence of forest species and the abundances of human-avoiding species and birds of prey; however, it favoured the abundances of forest resident hole-nesters and insectivores (Fig. 2). The proportion of water bodies in the landscape, usually within 500 m, 2 km, or 5 km scales, appears beneficial for most bird guilds, except for the functional richness of forest bird species (Fig. 2). Finally, the proportion of wetland areas decreased the abundances of forest resident hole-nesters, insectivores, and herbivores and the RaoQ of all species, while it promoted the functional divergence of all species and the RaoQ of forest species (Fig. 2).

The overall proportion of old forests promoted the abundance of human-avoiding species (5 km scale), presence of red-listed forest species (100 m scale), functional RaoQ of forest species (2 km scale), and the CWM of specialization index (5 km scale; Fig. 2). In contrast, the old forest proportion reduced the abundance of open-habitat species (100 m scale) but increased it at the 5 km scale (Fig. 2). The overall proportion of broadleaf forests promoted several guilds: the abundances of all species, open-habitat species (including the red-listed ones), human-avoiding species, birds of prey, and insectivorous species. Mixed forest proportion favoured the abundances of forest species, old-growth forest specialists, and herbivores at 100 m scale (Fig. 2) and increased functional diversity of all species while decreasing their functional divergence and RaoQ at the 500 m scale (Fig. 2).

## 4. Discussion

### 4.1. Finnish bird biodiversity: between energy, heterogeneity, and habitat amount

Our results showed that variations in Finnish breeding bird communities are due to a complex combination of energy constraints (Clarke & Gaston, 2006; Evans et al., 2005; Hawkins et al., 2003; Root, 1988) and habitat filtering via habitat heterogeneity and amount (Fahrig, 2013; Fahrig et al., 2011; Honkanen et al., 2010; Leibold et al., 2004).

#### 4.1.1. Energy primarily supports higher bird abundance but also species richness

As predicted by the *more-individual hypothesis*, latitude and soil fertility, as proxies for solar and geochemical energies, respectively, permitted higher abundances of overall bird communities and most species groups (Honkanen et al., 2010; Hurlbert, 2004; Luoto et al., 2007; Root, 1988). We found that higher average levels of tree volume, a proxy for energy availability, promoted the abundance of several guilds, as previously reported (Elo et al., 2012). Higher energy availability also supported the species richness of most species groups, although the effect was weaker than that for abundance. Since our models for species richness included abundance as a covariable, the latitude effects suggest an additional mechanism in the more-individual process, potentially the *diversification rate hypothesis* (Evans et al., 2005; Honkanen et al., 2010). Our results do not confirm that higher energy availability promotes more complex trophic chains (Evans et al., 2005). Our study shows diverging results manifested by two contrasting patterns: insectivores decreased northwards, as predicted by the hypothesis, probably due to lower arthropod abundance, while the abundance of birds of prey increased. However, we did not analyse trophic chains per se (i.e. birds themselves but not their preys and predators).

#### 4.1.2. Habitat heterogeneity positively affects both species diversity and abundance

We found mixed evidence supporting our hypothesis that habitat heterogeneity primarily affects species richness. Landscape heterogeneity, measured via the RaoQ of all habitats, promoted the species richness of several guilds and was always ranked high in metric importance within species richness models (Fig. S14), validating the hypothesis. Therefore, higher landscape habitat diversity seems to foster the diversity of available niches, directly enriching the number of species possibly present in a landscape (*sensu* the habitat-diversity hypothesis; Duflot et al., 2022; MacArthur & MacArthur, 1961). The importance of the RaoQ metric of all habitats increased in the models of more specialized guilds and had the strongest effects on the species richness of old-growth forest species (Fig. S14). In contrast, the Shannon diversity of all land cover and that of seminatural vegetation primarily affected abundance. The reason might be that rarer land covers, such as urban areas, cropland, and deciduous or mixed forests, shelter the highest avian abundance in Finland (see Fig. S15).

#### 4.1.3. Habitat amount primarily supports abundance of related guilds

Habitat amount was an important factor driving the abundance of the associated bird guilds, thereby confirming the *more-individual hypothesis*. The abundance of open-habitat species significantly increased with larger cropland proportions. We found that the overall forest proportion promoted the abundance and richness of forest bird species. Consequently, a larger proportion of forests might directly promote abundance via resource quantity and species richness via a potentially larger colonization–extinction ratio and greater habitat diversity (Fahrig, 2013; MacArthur & Wilson, 1967; Wright, 1983). Among the different forest habitats, old-growth and broadleaved forests were particularly important for forest birds, including red-listed ones, and human-avoiding species. In addition, the configuration of old-growth forest patches affected species richness, as large old forest patches with substantial core areas promoted the richness of forest and old-growth forest specialist species. Our results suggest that while habitat amount directly affects abundance, as predicted by the *more-individual hypothesis*, more complex processes are involved. For instance, large old forest patches combine interior habitats and edges, thereby increasing niche diversity and supporting bird species in both habitats (Barbaro et al., 2005; Terraube et al., 2016). In addition, broadleaved forests favoured open-habitat species, suggesting complementation processes in which they require resources provided by broadleaved trees (e.g. for nesting; Bosco et al., 2024; Virkkala et al., 2004).

### 4.2. Highly anthropic land uses negatively affect multiple bird guilds

Highly anthropic land uses (cropland and urban areas) showed negative effects on old-growth forest specialists, red-listed forest species, and human-avoiding species and led to an overall decrease in functional diversity. Intensification of farming practices is a known threat to many bird species in Finland and elsewhere, as highlighted in the Red List of Finnish Species (Lehikoinen et al., 2019). Our results show that disturbances from highly anthropic land uses (noises, light and chemical pollution, and human presence) are particularly detrimental to the conservation capacities of nearby forest habitat patches, with important edge effects in their surroundings (Barbaro et al., 2023; Fröhlich et al., 2025). However, forest resident hole-nesting species, which declined significantly in Fennoscandia due to forestry activities (Imbeau et al., 2001), seem to benefit from the conditions offered by urban areas. The installation of thousands of artificial cavity boxes near cities and houses in Finland might explain this positive result (Imbeau et al., 2001). These results advocate setting up larger protected areas to preserve typical bird communities of forest interiors (Fahrig, 2013). The positive effect of the mean patch size of old forests on the richness of forests and old-growth forest specialist species corroborates the need for large forest patches.

### 4.3. Higher latitudes fostered threatened species, specialization, and functional diversity

We found opposite latitudinal patterns between taxonomic and functional diversity. Most taxonomic variables decreased northwards (see 4.1), while the overall functional diversity of all species and forest species increased northwards (Fig. 3). In addition, community specialization and the presence of red-listed forest species increased northwards (Fig. 3). Under the *energy-diversity hypothesis*, increasing energy should support additional rare resources and, consequently, specialist species (*energy-specialization hypothesis*; Evans et al., 2005). Our results contradict this assumption, since higher energy translated into a greater abundance of functionally similar species, while lower energy led to communities composed of functionally dissimilar species. Bae *et al*. (2018) found similar results to ours in temperate forests by measuring energy as productivity using the normalized difference vegetation index (NDVI). In boreal and temperate forest landscapes, less productive sites have historically been less harvested than productive ones, thereby allowing for rare habitats and associated specialized species to persist (Mönkkönen et al., 2022). In Finland, this may explain the higher proportion of red-listed forest bird species northwards (Fig. 3).

Boreal species are less competitive but more specialized than their temperate relatives; in the set of species we considered, we found a significant negative relationship between species specialization and temperature indices (*p* < 0.001). Therefore, our findings might reflect the competitive exclusion of the more specialized cold-dwelling species in southern latitudes and warm-dwelling generalist species in high latitudes (Callaghan et al., 2004; Cours et al., 2025). A similar pattern of community specialization along the latitudinal gradients was found at the European scale, suggesting that extreme northern climatic conditions have resulted in selected specialized bird species (Rivas-Salvador et al., 2019). Faster warming at higher latitudes compared to southern areas (Loarie et al., 2009) might be another reason for the increased threat to northern bird communities (Mönkkönen et al., 2022; Virkkala et al., 2008; Virkkala & Rajasärkkä, 2010). Such specialized, threatened, and functionally diverse communities are particularly vulnerable and therefore require particular conservation attention (Santangeli et al., 2017).

### 4.4. Biodiversity conservation: a matter of specialized landscapes

Anthropogenic land use activities strongly decreased the presence of threatened forest species, as well as the old-growth forest specialists, and human-avoiding species, with effects observed for up to 5 km. Therefore, forest protection areas located far from human land uses, such as cropland or urban areas, and embedded within low-intensity forestry matrices are likely to achieve conservation goals (Brazner et al., 2024; Fröhlich et al., 2025; Häkkilä et al., 2017). In contrast, cropland and urban areas promoted the abundance of open-habitat species, including threatened ones. Appropriate management intensity and practices should be adopted to maintain the biodiversity of such landscapes. Finally, landscape habitat heterogeneity increased biodiversity variables, strengthening the previous evidence (Cours & Duflot, 2025; Duflot et al., 2022; Fahrig et al., 2011; MacArthur & MacArthur, 1961) that it is an important component of biodiversity. We suggest that a combination of diverse landscapes with high, moderate, and minimum human impact should be preserved to support the overall diversity of bird species at regional scales (Duflot et al., 2014).

## Acknowledgements

The authors are very grateful and thankful for the many birdwatchers who made the present study (and many others) possible.

