## Supplementary Information for "Energy and heterogeneity shape bird taxonomic and functional gamma-diversity patterns across landscapes in Finland"

### 1. Detailed methodology of the line transects surveys

The line transect method is a one-visit census, in which birds are identified and counted along circa 6 km-long transects (in our analysis: mean = 6,056 m, min = 2,999 m & max = 7,299 m). These transects are systematically distributed across Finland on a grid of 25 km ´ 25 km and are designed and positioned in advance to account for general region habitat composition (i.e. forests, mires, croplands, etc.; Järvinen, Koskimies and Väisänen, 1991; Virkkala and Lehikoinen, 2014). A sample of line transects are visited each year by an observer counting pairs of birds either within 25 meters on both sides of the transect line, namely the "main belt", or beyond 25 meters, as far as birds can be observed, namely the "supplementary belt". Both belts combined form the "survey belt". The visits are typically made from late May to late June based on geographical position, bird breeding season starting earlier in Southern Finland. Visits are performed in the early mornings (usually between 3:00 and 9:00 pm), at the peak singing activity, in dry weather conditions.

We calculated species richness and abundance (number of breeding pairs) based on both the main and the supplementary belt – i.e. within the survey belt. To account for differences in species detectability between the main and the supplementary belt, we calculated species-specific correction coefficients based on distance of sampling (Virkkala and Lehikoinen, 2014). The value of the coefficient is low for species frequently observed within the supplementary belt (to avoid potential over-representation of these species due to high relative long-distance detectability) and is high for species mostly observed within the main belt. We calculated this coefficient based on observations done in all available transects (i.e. from 2006 to 2022) across the entire Finland; the larger the data the more reliable the coefficients are (Virkkala and Lehikoinen, 2014). We applied the correction coefficient on the abundance of each species observed in the survey belt, rounded up to the nearest integer to keep the characteristic of count data for statistical models.

Since we focused on terrestrial ecosystems, we only included in our analysis the species inhabiting forest, woodland, shrubland, wetland, grassland, riverine, rock, and human modified habitats, therefore excluding coastal and marine species (source for habitat preferences: Tobias *et al.* (2022). We also removed Strigidae since their observations are not reliable using the transect census method (Virkkala and Lehikoinen, 2014).

Järvinen, O., Koskimies, P. and Väisänen, R.A. (1991) "Line Transect Census of BreedingLand Birds," *Monitoring Bird Populations. A Manual of Methods Applied in Finland*. Zoological Museum, Finnish Museum of Natural History. Helsinki, Finland, pp. 33–41. Available at: https://tietopankki.luomus.fi/wp-content/uploads/2024/04/03-Line-transect-census.pdf (Accessed: May 15, 2025).

Tobias, J.A. *et al.* (2022) "AVONET: morphological, ecological and geographical data for all birds," *Ecology Letters*, 25(3), pp. 581–597. Available at: https://doi.org/10.1111/ele.13898.

Virkkala, R. and Lehikoinen, A. (2014) "Patterns of climate-induced density shifts of species: poleward shifts faster in northern boreal birds than in southern birds," *Global Change Biology*, 20(10), pp. 2995–3003. Available at: https://doi.org/10.1111/gcb.12573.

#### 2. Corine Land Cover broad classes correspondence

**Table S1** | Correspondence between raw values from Corine Land Cover raster data and our broad land cover classes.

| Original raster value | Broad land cover classes |
|---|---|
| **Corine Land Cover 2006** | |
| 1-13 | Urban areas |
| 14-16 | Cropland |
| 17, 26, 27 | Grass & Moorland |
| 18-25 | Forest |
| 28-33 | Woodland & Scrubland |
| 34, 35 | Sand, bare & rock |
| 36-41 | Wetland |
| 42-44 | Water |
| **Corine Land Cover 2012** | |
| 1-15 | Urban areas |
| 16, 17, 20, 21 | Cropland |
| 18, 19, 30, 31 | Grass & Moorland |
| 22-29 | Forest |
| 32-36 | Woodland & Scrubland |
| 37-39 | Sand, bare & rock |
| 40-45 | Wetland |
| 46-48 | Water |
| **Corine Land Cover 2018** | |
| 1-16 | Urban areas |
| 17, 18, 21, 22 | Cropland |
| 19, 20, 31, 32 | Grass & Moorland |
| 23-30 | Forest |
| 33-37 | Woodland & Scrubland |
| 38-40 | Sand, bare & rock |
| 41-46 | Wetland |
| 47-49 | Water |

#### 3. Discrete classes for MS-NFI continuous data

**Table S2** | Discrete classes used to simplified MS-NFI continuous data

| Discrete classes | Continuous values |
|---|---|
| **Canopy cover** | |
| C0 | 0 |
| C0_5 | ]0 ; 5[ |
| C5_20 | [5 ; 20[ |
| C20_40 | [20 ; 40[ |
| C40_60 | [40 ; 60[ |
| C60_80 | [60 ; 80[ |
| C80_100 | [80 ; 100] |
| **Diameter at breast height** | |
| D0 | 0 |
| D0_10 | ]0 ; 10[ |
| D10_30 | [10 ; 30[ |
| D30_40 | [30 ; 40[ |
| D40_more | [40 ; more] |
| **Tree height** | |
| H0 | 0 |
| H0_1 | ]0 ; 1[ |
| H1_5 | [1 ; 5[ |
| H5_10 | [5 ; 10[ |
| H10_20 | [10 ; 20[ |
| H20_30 | [20 ; 30[ |
| H30_more | [30 ; more] |
| **Tree age** | |
| A0 | 0 |
| A0_20 | ]0 ; 20[ |
| A20_40 | [20 ; 40[ |
| A40_60 | [40 ; 60[ |
| A60_80 | [60 ; 80[ |
| A80_100 | [80 ; 100[ |
| A100_120 | [100 ; 120[ |
| A120_140 | [120 ; 140[ |
| A140_more | [140 ; more] |
| **Tree total volume** | |
| V0 | 0 |
| V0_10 | ]0 ; 10[ |
| V10_30 | [10 ; 30[ |
| V30_60 | [30 ; 60[ |
| V60_90 | [60 ; 90[ |
| V90_120 | [90 ; 120[ |
| V120_150 | [120 ; 150[ |
| V150_more | [150 ; more] |
| **Broadleaf tree volume** | |
| Vbroad0 | 0 |
| Vbroad0_10 | ]0 ; 10[ |
| Vbroad10_30 | [10 ; 30[ |
| Vbroad30_60 | [30 ; 60[ |
| Vbroad60_90 | [60 ; 90[ |
| Vbroad90_120 | [90 ; 120[ |
| Vbroad120_150 | [120 ; 150[ |
| Vbroad150_more | [150 ; more] |

#### 4. Extrapolated vegetation types

**Table S3** | Semi-natural vegetation types based on both CLC and MS-NFI data.

| Type of vegetation | Rules |
|---|---|
| Bare_soil | Volume tree = 0 & CLC = Sand, bare soil, rock |
| Herbaceous_Shrub | Volume tree = 0 & CLC = Grass & moorland, Wetland mires |
| Shrub | Volume tree = 0 & CLC = Forest, Woodland & Scrub, Wooded wetland |
| Pine | Volume tree > 0 & Pine ≥ 80% of Volume tree total |
| Spruce | Volume tree > 0 & Spruce ≥ 80% of Volume tree total |
| Birch | Volume tree > 0 & Birch ≥ 80% of Volume tree total |
| Other_broadleaf | Volume tree > 0 & Other broadleaf ≥ 80% of Volume tree total |
| Coniferous_dominant | Volume tree > 0 & Pine + Spruce ≥ 80% of Volume tree total |
| Broadleaf_dominant | Volume tree > 0 & Birch + Other broadleaf ≥ 80% of Volume tree total |
| Mixed_spp | Volume tree > 0 & Coniferous and Broadleaf < 80% of Volume tree total |

#### 5. Landscape environmental variables

**Table S4** | List of landscape variables selected for the study.

| Variable | Definition |
|---|---|
| Shannon CLC | Shannon heterogeneity index based on Corine Land Cover classes. |
| Shannon Semi nat | Shannon heterogeneity index based on extrapolated vegetation types (see Table S3) in semi-natural land cover classes (i.e. Forest, Woodland & Scrubland, Wetland, and Grass & Moorland, see Table S1). |
| FRic All | "Habitat species" functional richness in all land cover classes (see Figure S1). |
| RaoQ All | "Habitat species" functional RaoQ in all land cover classes (see Figure S1). |
| FDiv All | "Habitat species" functional divergence in all land cover classes (see Figure S1). |
| FRic Semi nat | "Habitat species" functional richness in semi-natural land cover classes (see Figure S1). |
| RaoQ Semi nat | "Habitat species" functional RaoQ in semi-natural land cover classes (see Figure S1). |
| FDiv Semi nat | "Habitat species" functional divergence in semi-natural land cover classes (see Figure S1). |
| CWM Canopy | Average canopy closure of the vegetation in forested areas. |
| CWM DBH | Average diameter at breast height of the vegetation in forested areas. |
| CWM Age | Average age of the vegetation in forested areas. |
| CWM Vol tot | Average total tree volume in forested areas. |
| CWM Vol broad | Average broadleaf tree volume in forested areas. |
| CWM Fertility | Average fertility level in semi-natural classes. |
| FDis Canopy | Functional dispersion of vegetation canopy closure in forested areas. |
| FDis DBH | Functional dispersion of vegetation diameter at breast height in forested areas. |
| FDis Age | Functional dispersion of vegetation age in forested areas. |
| FDis Vol tot | Functional dispersion of total tree volume in forested areas. |
| FDis Vol broad | Functional dispersion of broadleaf tree volume in forested areas. |

| | |
|---|---|
| FDis Fertility | Functional dispersion of fertility level in semi-natural classes. |
| Old forest amount | Total proportion of old forests (age ≥ 100 years). |
| Mn core area Old | Mean core area of all old forest patches (*landscapemetrics* R-package). |
| Mn para Old | Mean perimeter-area ratio of all old forest patches (*landscapemetrics* R-package). |
| Broad forest amount | Total proportion of broadleaf forests (volume broadleaf ≥ 80% total tree volume). |
| Mn core area Broad | Mean core area of all broadleaf forest patches (*landscapemetrics* R-package). |
| Mn para Broad | Mean perimeter-area ratio of all broadleaf forest patches (*landscapemetrics* R-package). |
| Mixed forest amount | Total proportion of mixed forests (volume broadleaf ≥ 80% total tree volume). |
| Mn core area Mixed | Mean core area of all mixed forest patches (*landscapemetrics* R-package). |
| Mn para Mixed | Mean perimeter-area ratio of all mixed forest patches (*landscapemetrics* R-package). |
| Edge dens Mixed | Edge density of all mixed forest patches (*landscapemetrics* R-package). |
| Patch dens Mixed | Density of all mixed forest patches (*landscapemetrics* R-package). |
| Mn core area Semi nat Plant | Mean core area of all semi-natural patches based on extrapolated vegetation types (*landscapemetrics* R-package). |
| Mn para Semi nat Plant | Mean perimeter-area ratio of all semi-natural patches based on extrapolated vegetation types (*landscapemetrics* R-package). |
| Mn patch dens Semi nat Plant | Density of semi-natural patches based on extrapolated vegetation types (*landscapemetrics* R-package). |
| Mn core area CLC | Mean core area of all Corine Land Cover patches (*landscapemetrics* R-package). |
| Mn para CLC | Mean perimeter-area ratio of all Corine Land Cover patches (*landscapemetrics* R-package). |
| Sum clearcut 5yr | Cumulated proportion of forest clearcut in landscape over the 5 years preceding bird observation. |
| Mean patch area clearcut 5yr | Mean area of forest clearcut patches cumulated over the 5 years preceding bird observation. |
| Forest prop. | Total forest proportion in buffer. Pixels of Forest, Woodland & Scrubland, and Wetland having canopy closure > 20%. |
| Cropland prop. | Total cropland proportion in buffer. |
| Urban prop. | Total Urban area proportion in buffer. |
| Water prop. | Total water (mostly lakes) proportion in buffer. |

#### 6. Calculation functional landscape heterogeneity

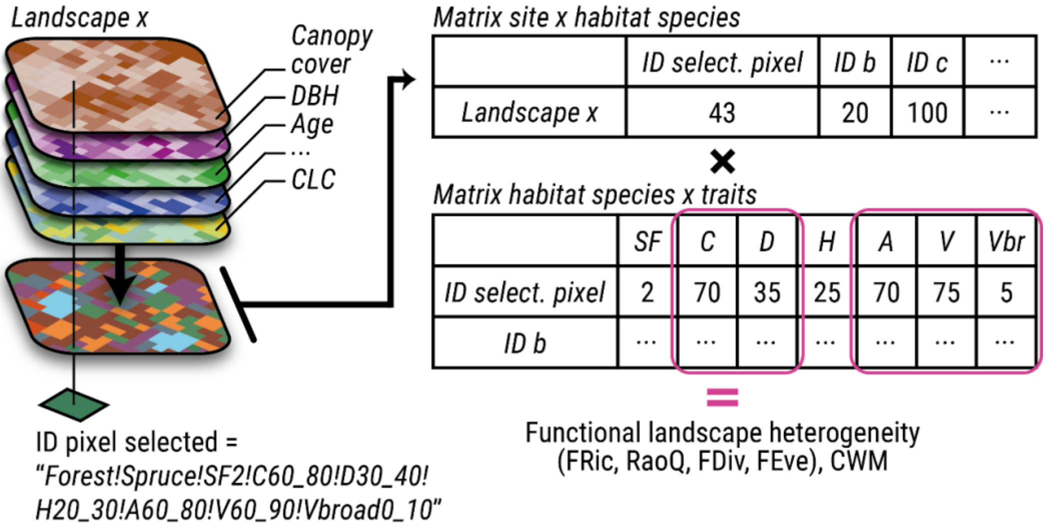


**Fig. S1** | Calculation method for the functional landscape heterogeneity. We use the landscape as a sampling area populated by "habitat species" gathering multisource information. We therefore use that information as traits related to each species, traits that we can use in addition to habitat species abundance to calculate functional index, but also community weighted mean, trait's dispersion or amount of specific habitats. "*SF*" = "*Site fertility*", "*C*" = "*Canopy cover*", "*D*" = "*Diameter at breast height*", "*H*" = "*Tree height*", "*A*" = "*Tree age*", "*V*" = "*Volume total*", and "*Vbr*" = "*Volume broadleaf*". Only canopy cover, tree diameter, age, total volume, and broadleaf volume were used for the calculation of Fric, RaoQ, FDiv, and FEve.

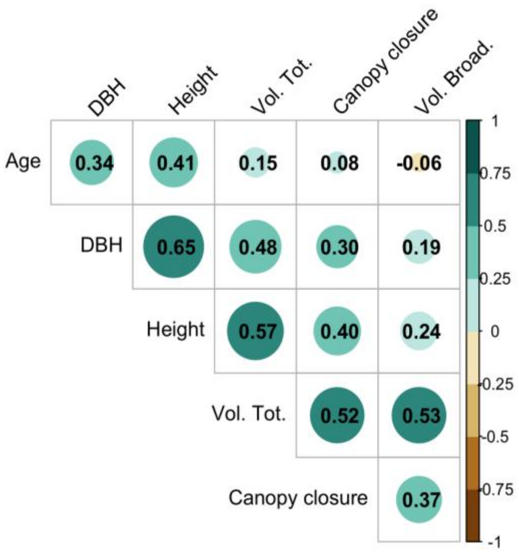


**Fig. S2** | Correlations of "habitat species" traits.

#### 7. Correlations between predictors

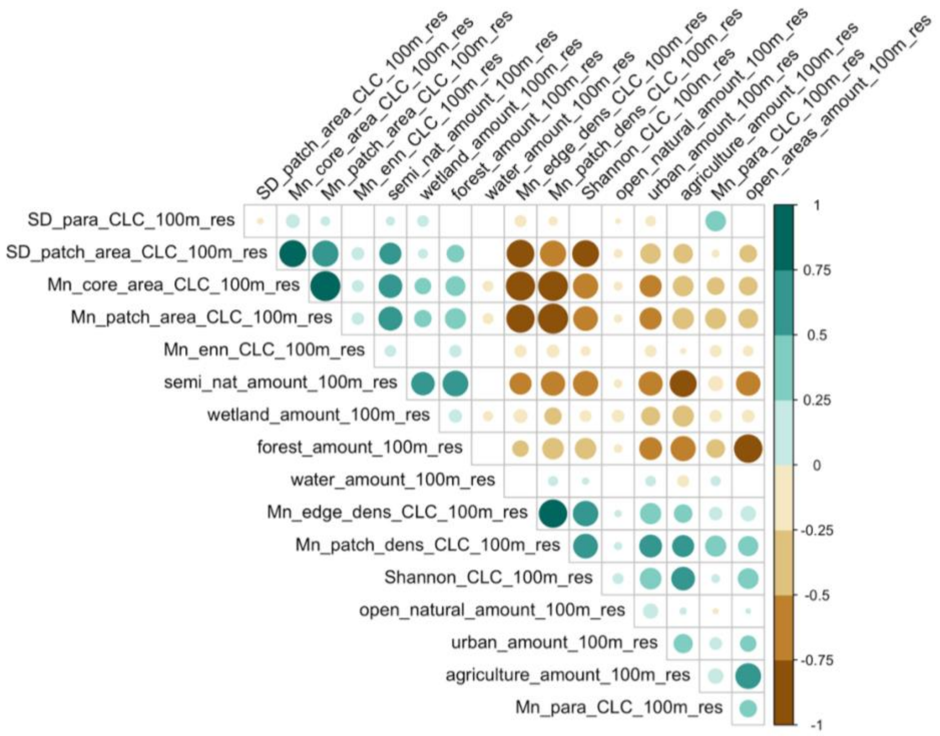


**Fig. S3** | Spearman correlation between Land Covers predictors at 100m.

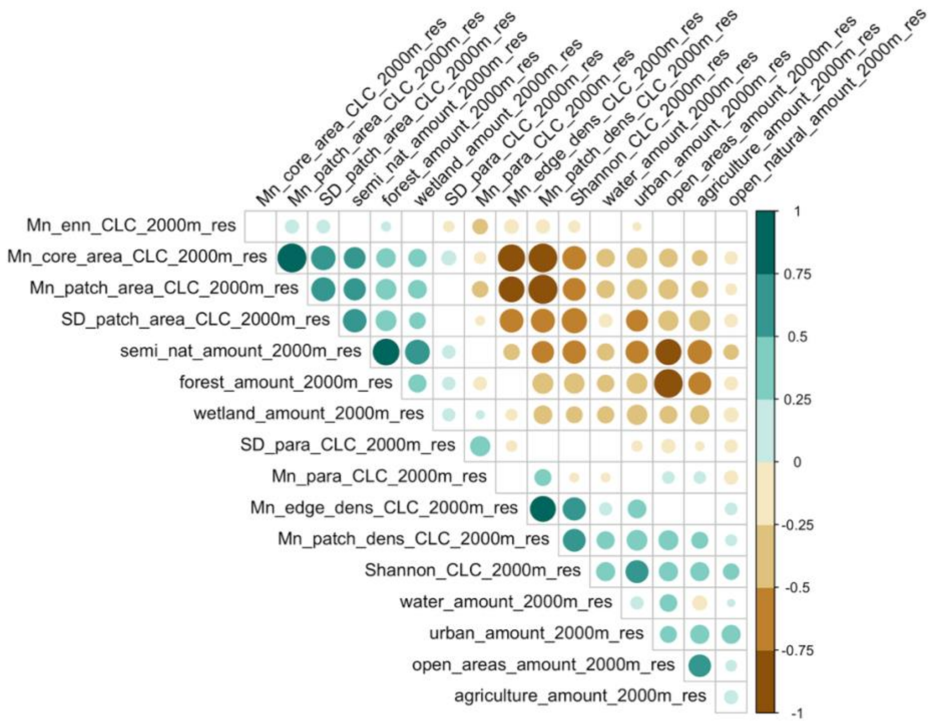


**Fig. S4** | Spearman correlation between Land Covers predictors at 2000m.

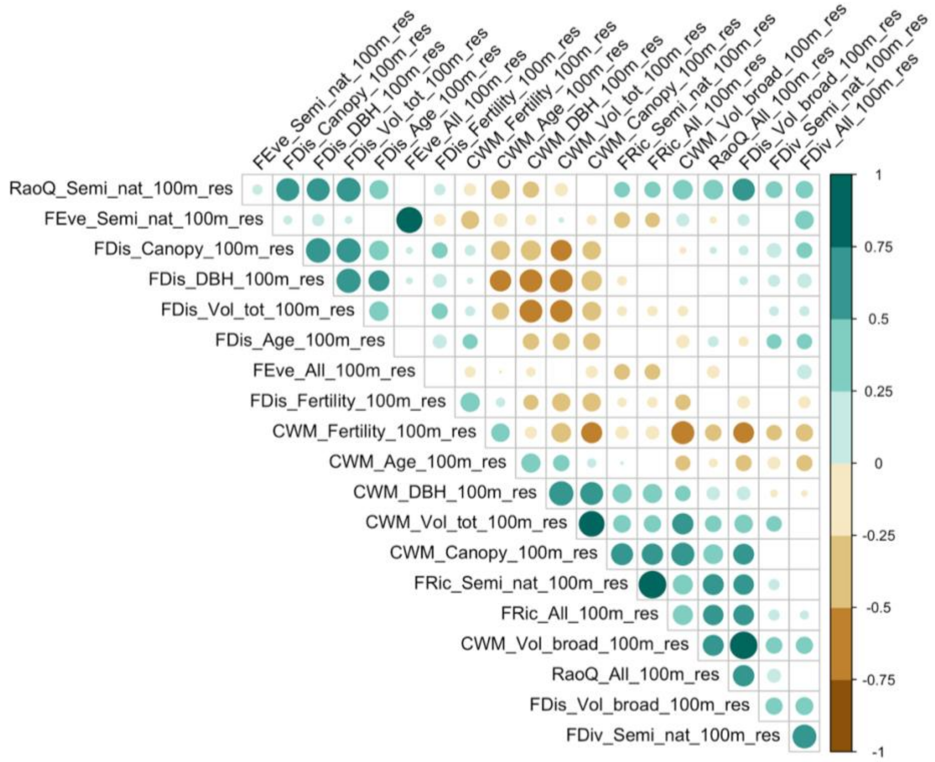


**Fig. S5** | Spearman correlation between landscape functional predictors at 100m.

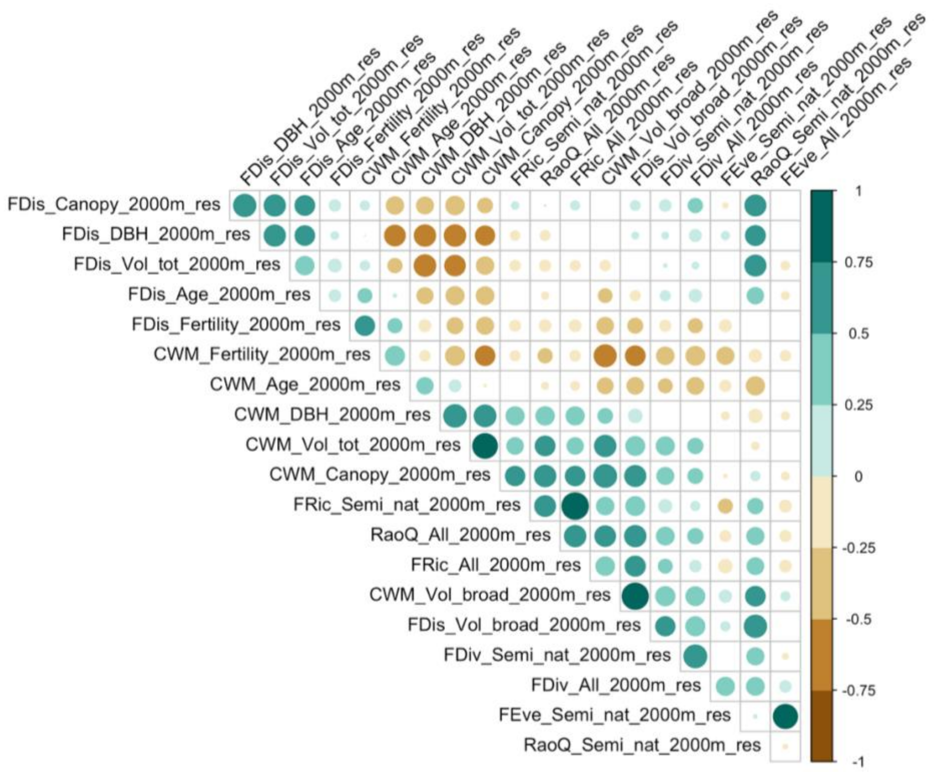


**Fig. S6** | Spearman correlation between landscape functional predictors at 2000m.

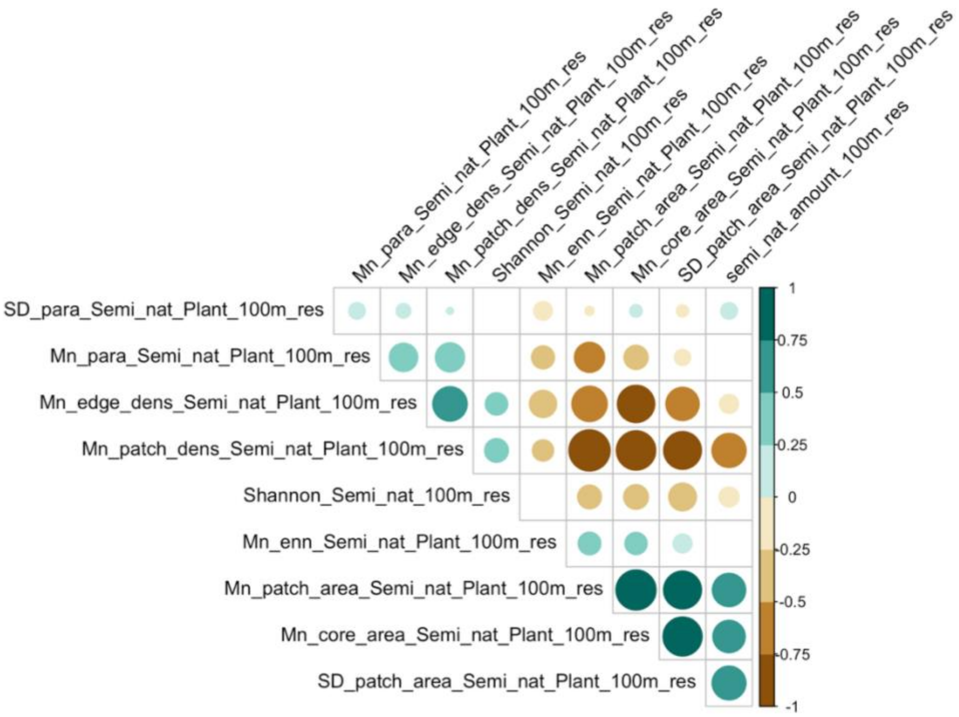


**Fig. S7** | Spearman correlation between Semi-Natural predictors at 100m.

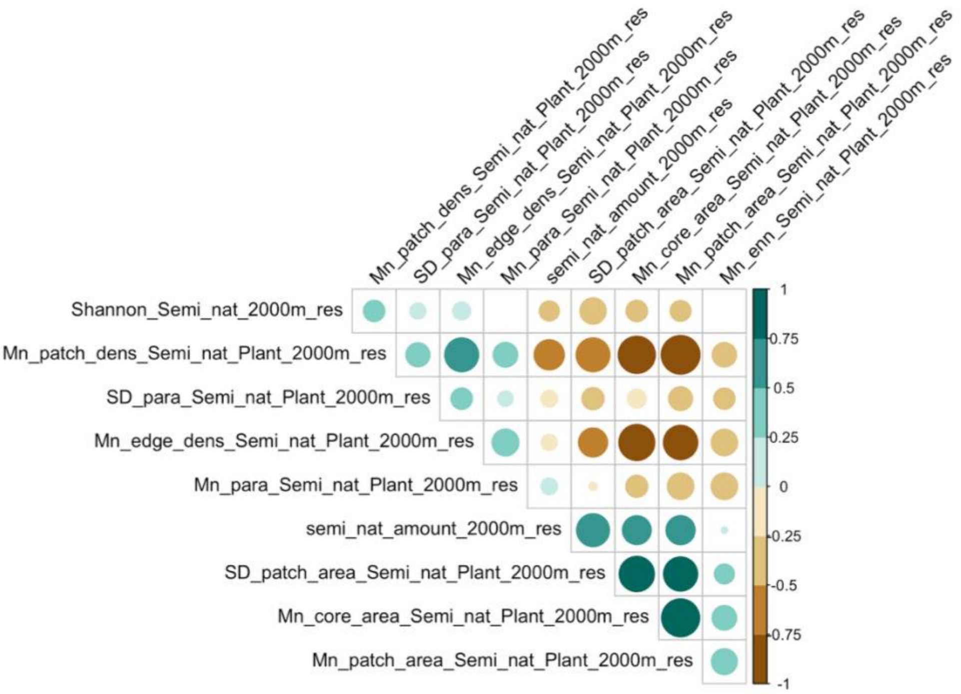


**Fig. S8** | Spearman correlation between landscape functional predictors at 2000m.

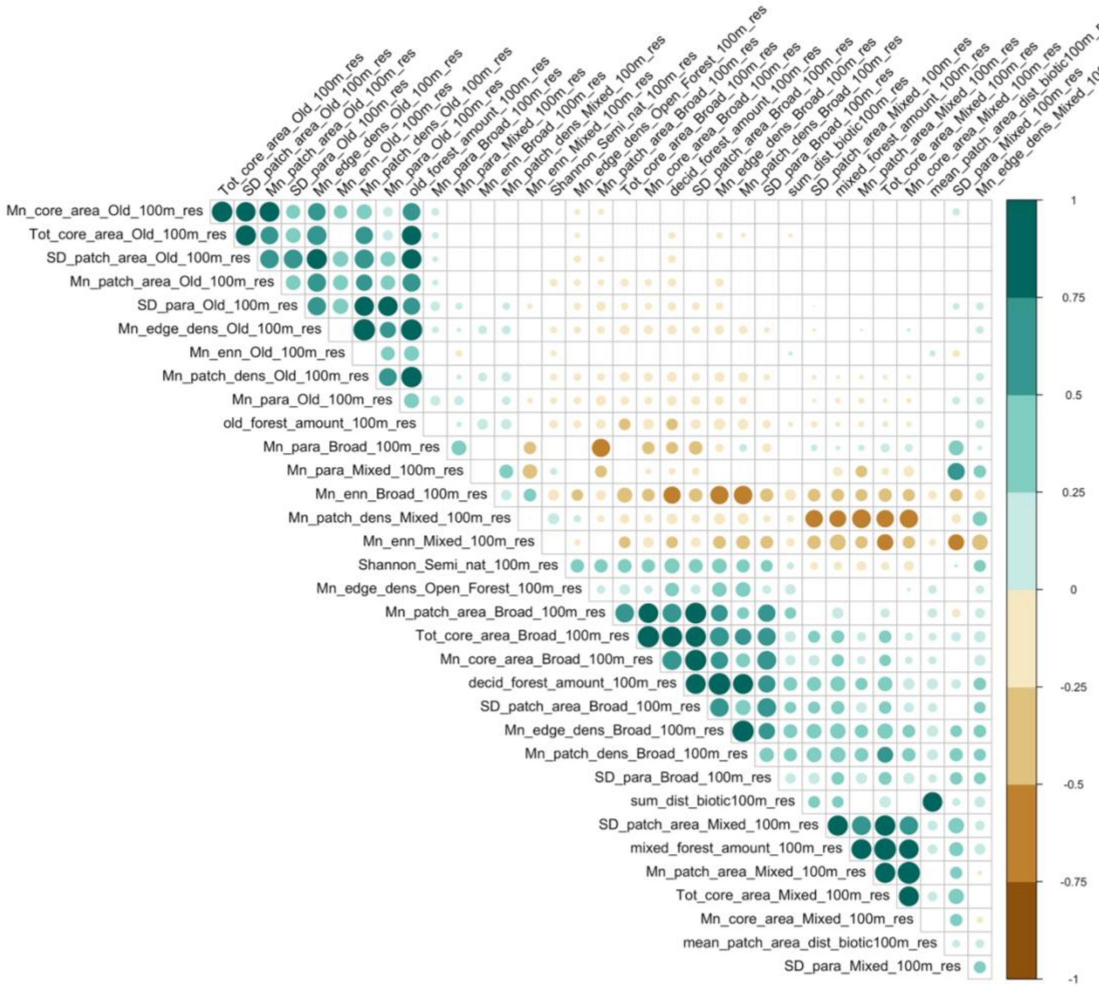


**Fig. S9** | Spearman correlation between Forest predictors at 100m.

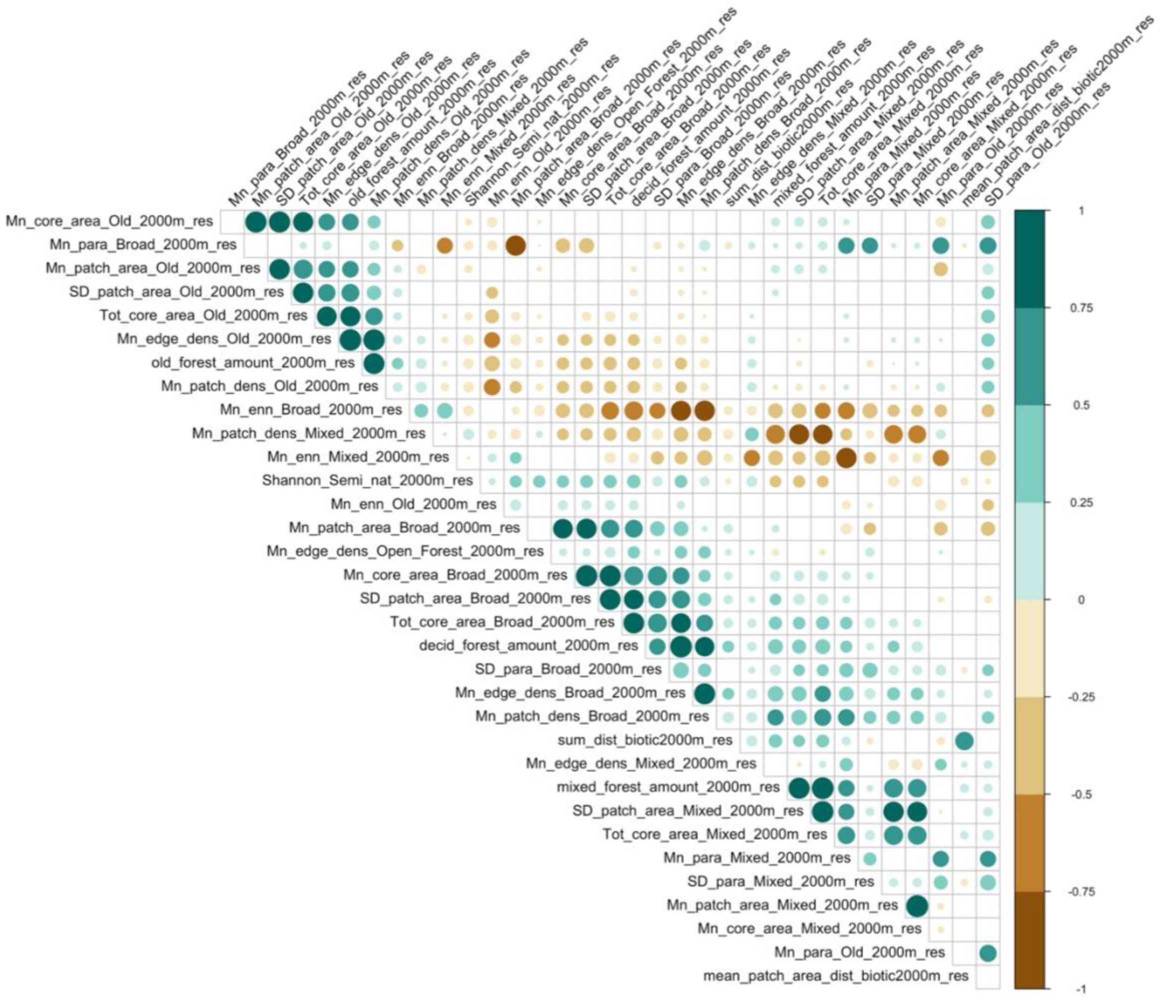


**Fig. S10** | Spearman correlation between Forest predictors at 2000m.

#### 8. Biodiversity metric presentation

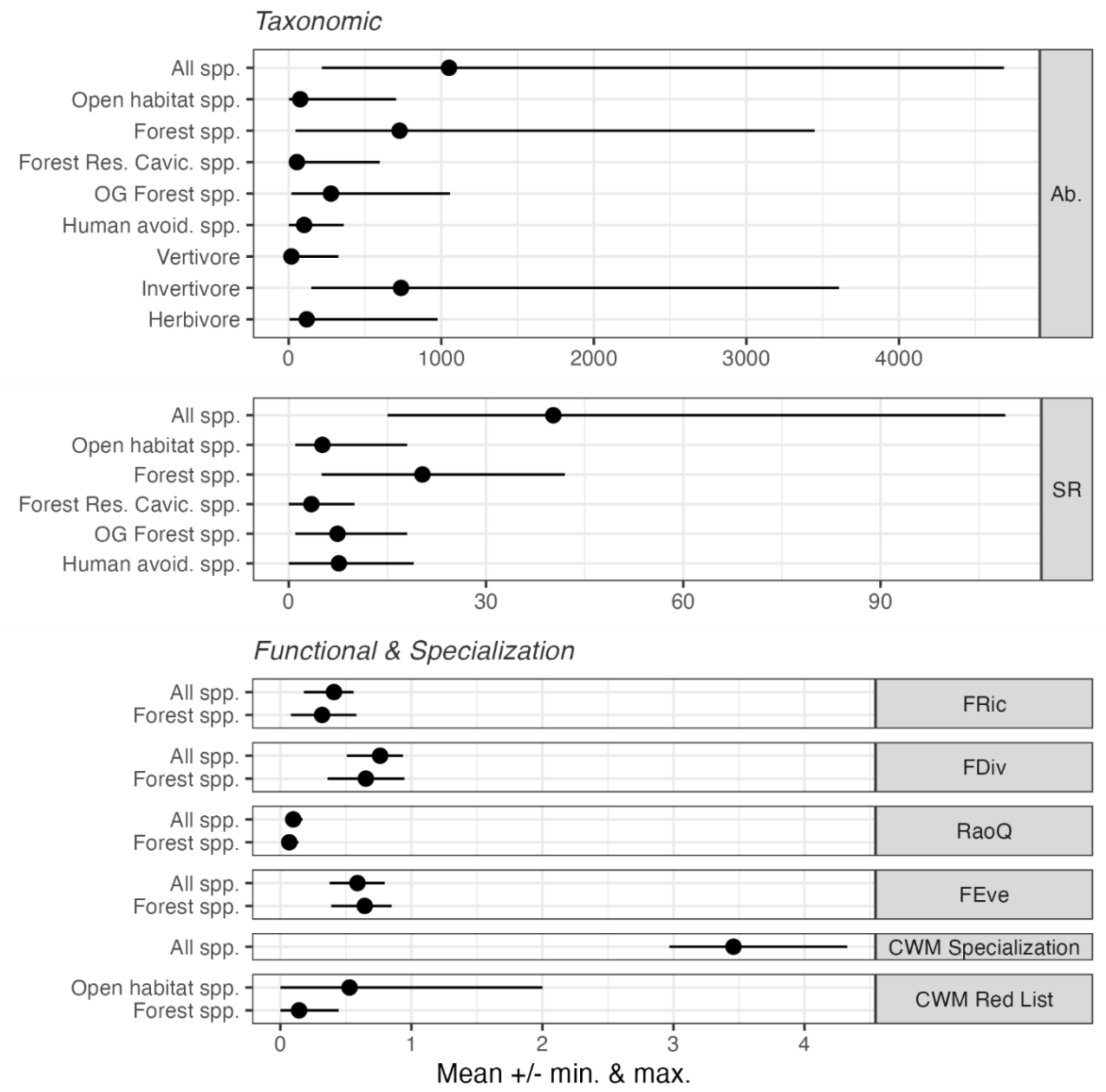


**Fig. S11** | Mean and range (minimum and maximum) values for each of the biodiversity variables presented in the study.

### 9. Spatial autocorrelation

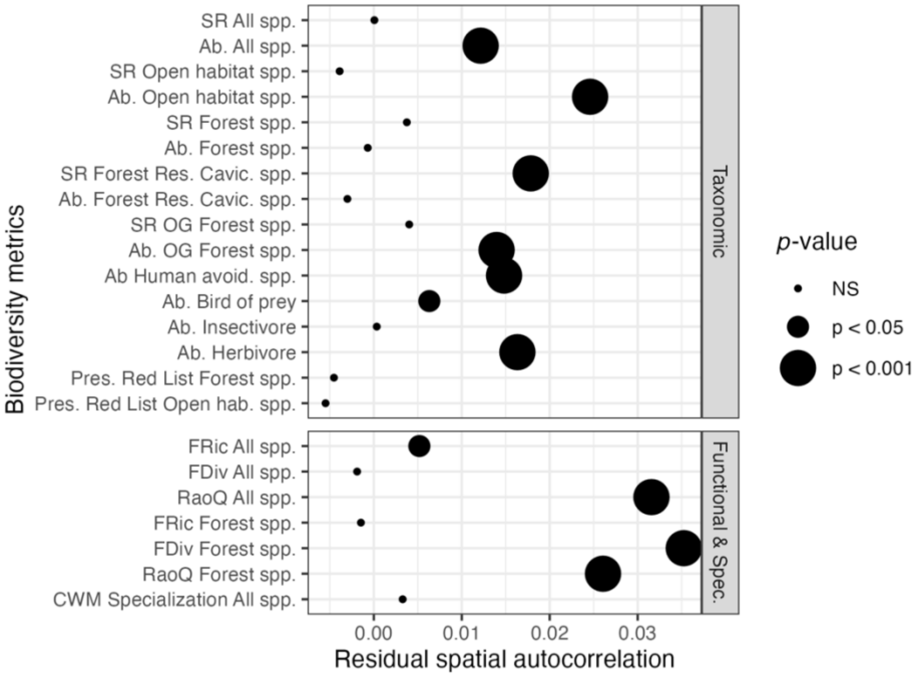


**Fig. S12** | Spatial autocorrelation for each biodiversity metric model.

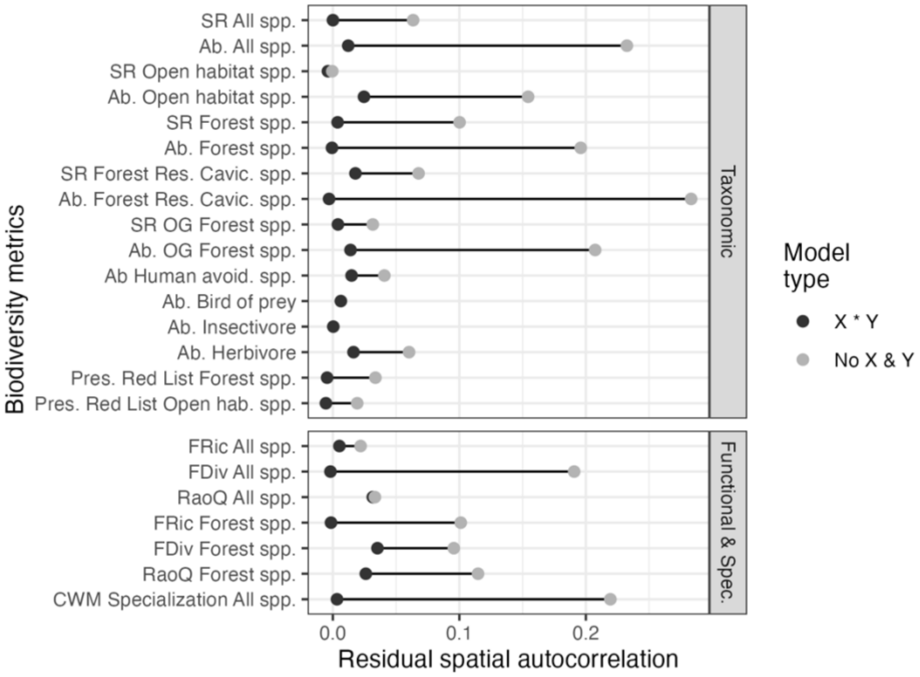


**Fig. S13** | Spatial autocorrelation for each biodiversity metric model. In black, autocorrelation from models including both latitude and longitude, and their interaction. In grey, models without them. While significant in many models including latitudes and longitudes (Fig. S4), values of autocorrelation remain very low compared to model not including geographical information.

#### 10. Predictors importance

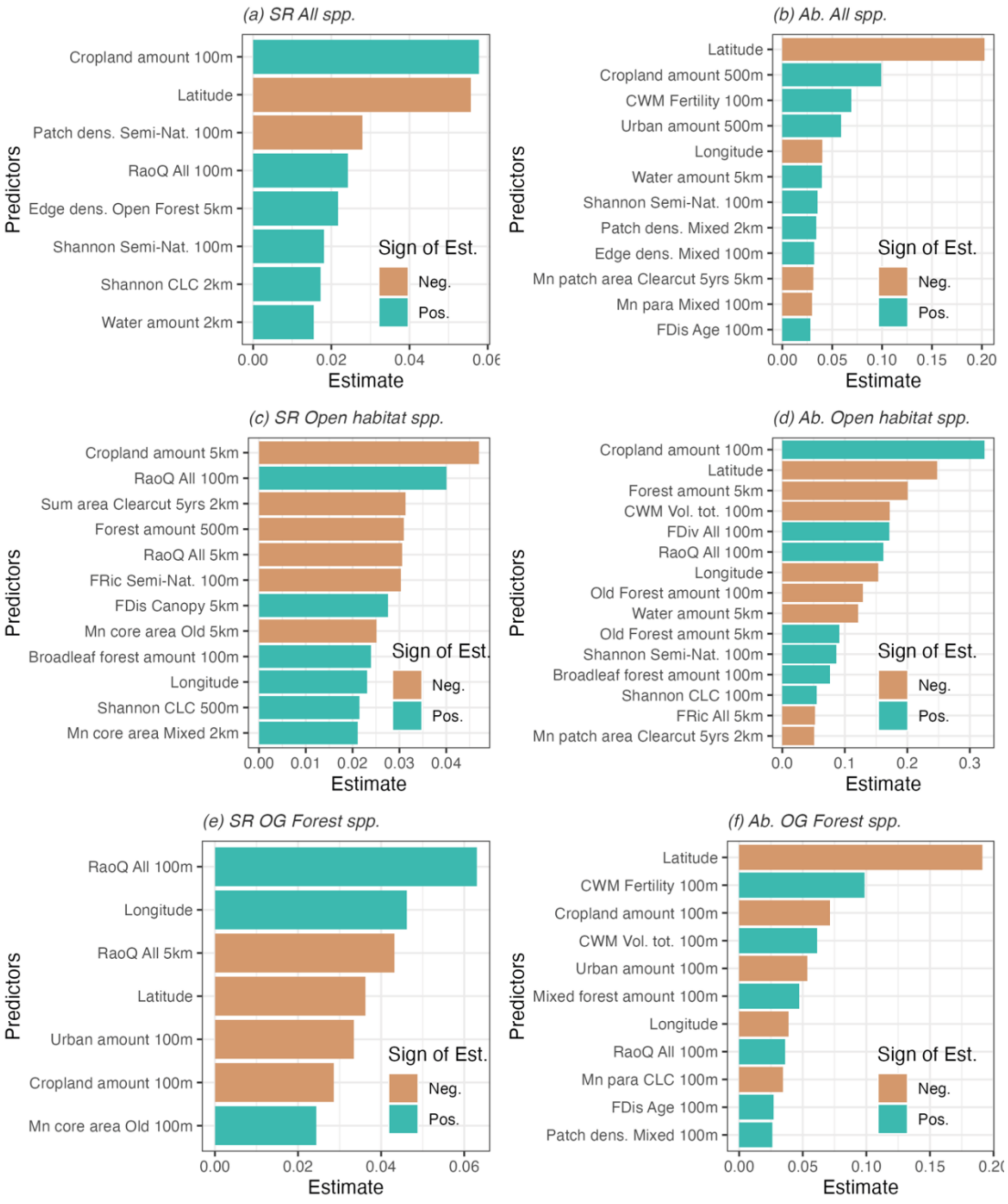


**Fig. S14** | Landscape metric relative importance in models, based on their estimates, for a selection of biodiversity metrics. Sign of the Estimate colour code indicate positive and negative effects.

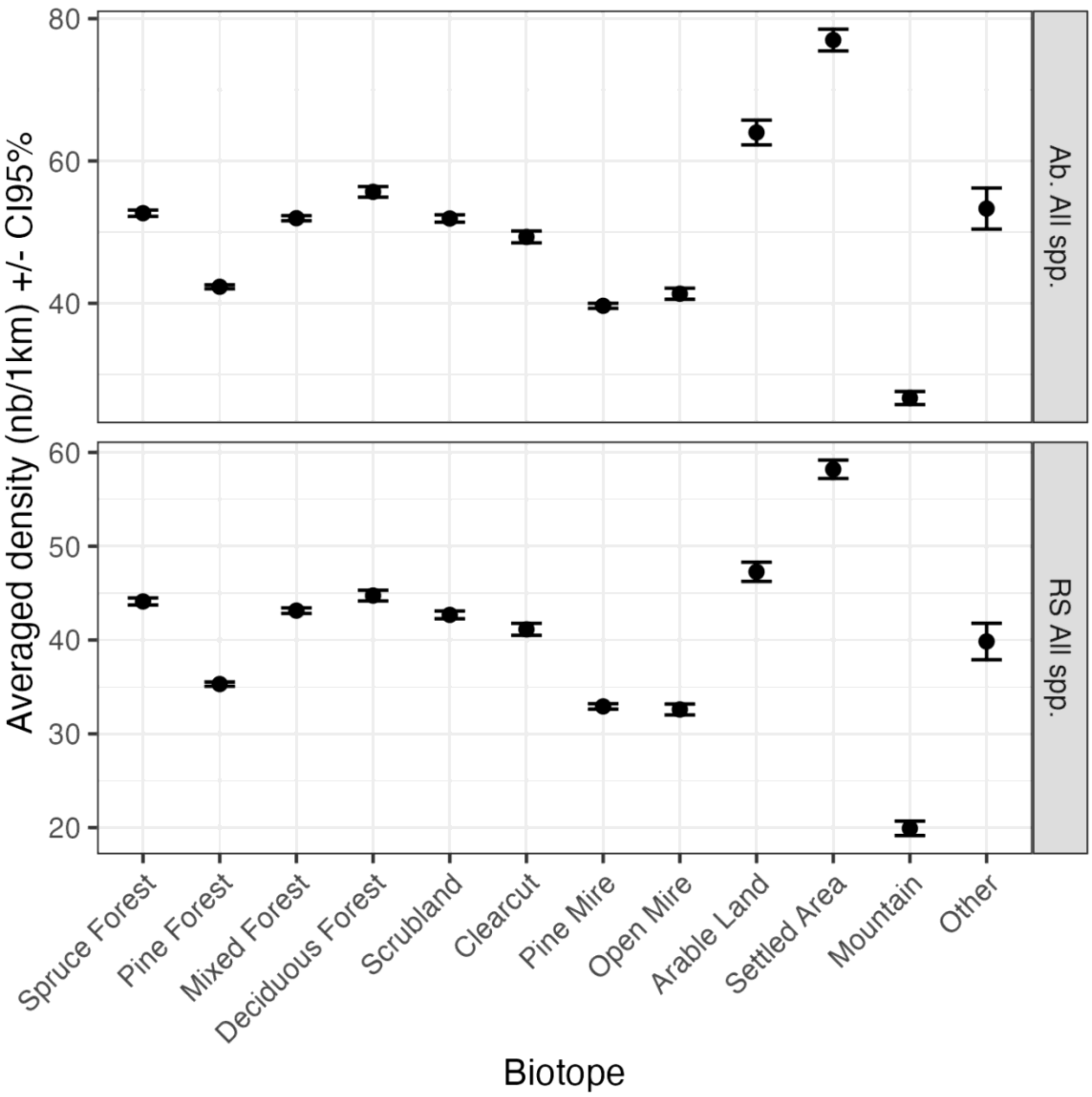


**Fig. S15** | Abundance (top) and species richness (bottom) of all bird species at the local (alpha)-scale. The density is calculated based on information regarding each section of bird monitoring transect, directly informed by bird observers.

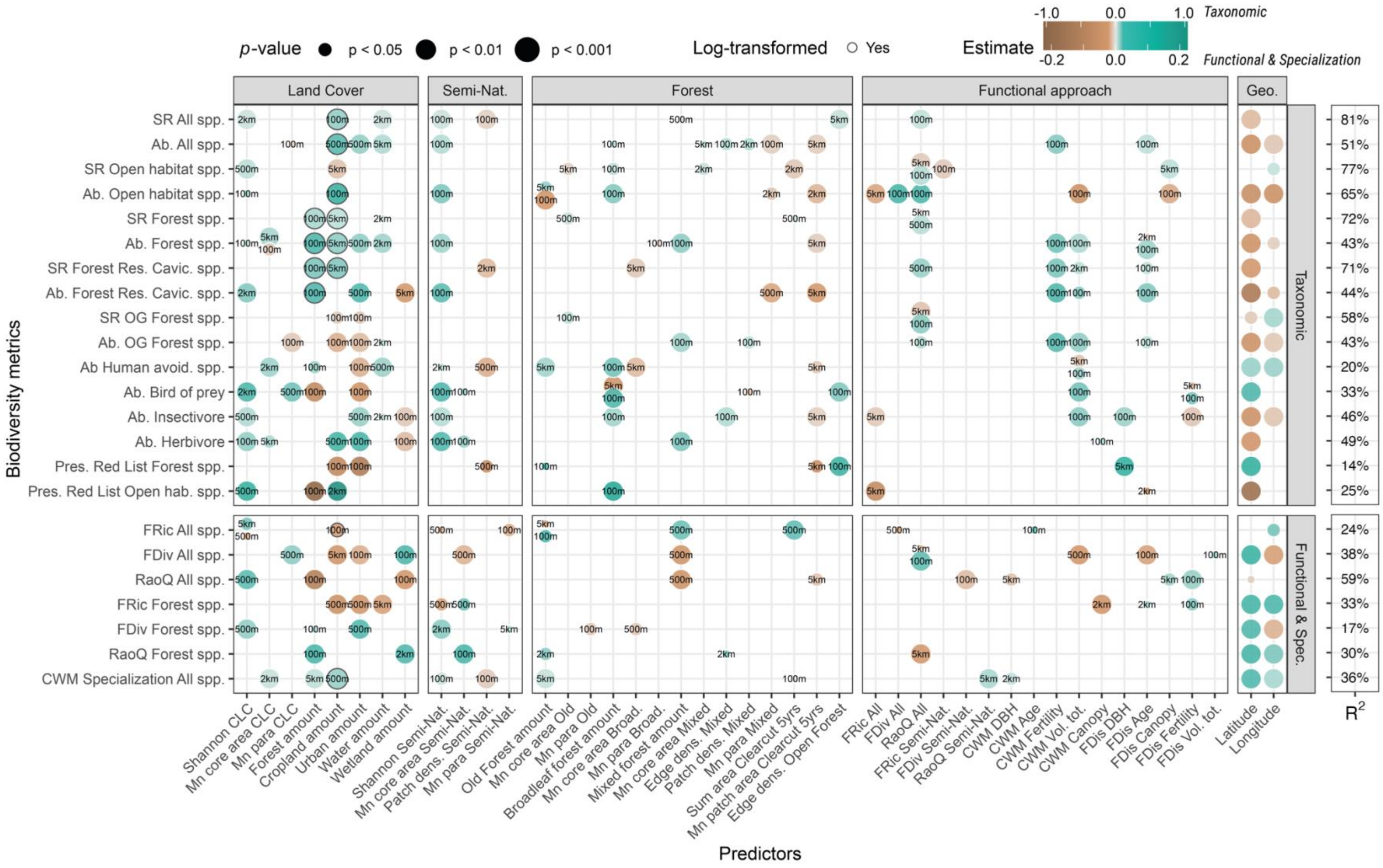


**Fig. S16** | Graphical representation of significant landscape metrics effects on bird taxonomic diversity (top) and functional and specialization indices (bottom). Number in bubbles indicates the landscape scale at which the effects were detected. R-squared are the explained deviance in models.
